# Pavlovian fear conditioning does not readily occur in rats in naturalistic environments

**DOI:** 10.1101/2021.10.20.465116

**Authors:** Peter R. Zambetti, Bryan P. Schuessler, Bryce E. Lecamp, Andrew Shin, Eun Joo Kim, Jeansok J. Kim

**Affiliations:** Department of Psychology, University of Washington, Seattle, WA 98195; Undergraduate Program in Neuroscience, University of Washington, Seattle, WA 98195; Undergraduate Program in Human Biology, Stanford University, Stanford, CA 94305

**Keywords:** Learning, memory, ethology, anxiety, PTSD, sensitization

## Abstract

Pavlovian fear conditioning, which offers the advantage of simplicity in both the control of conditioned and unconditioned stimuli (CS, US) presentation and the analysis of specific conditioned and unconditioned responses (CR, UR) in a controlled laboratory setting, has been the standard model in basic and translational fear research. Despite 100 years of experiments, the utility of fear conditioning has not been trans-situationally validated in real-life contexts. We thus investigated whether fear conditioning readily occurs and guides the animal’s future behavior in an ecologically-relevant environment. To do so, Long-Evans rats foraging for food in an open arena were presented with a tone CS paired with electric shock US to their dorsal neck/body that instinctively elicited escape UR to the safe nest. On subsequent test days, the tone-shock paired animals failed to exhibit fear CR to the CS. In contrast, animals that encountered a realistic agent of danger (a looming artificial owl) paired with a shock, simulating a realistic predatory strike, instantly fled to the nest when presented with a tone for the first time. These results illustrate the survival function and precedence of a nonassociative process, rather than associative conditioning, in life-threatening situations that animals are likely to encounter in nature.

## Introduction

Since the time of Watson and Morgan’s (1) conception that emotions, such as fear, should be studied as conditioned (acquired) reactions and Watson and Rayner’s (2) demonstration that fear can be rapidly learned in 9-month-old “Little Albert,” Pavlovian (or classical) fear conditioning has been the paradigm par excellence for studying both normal and abnormal fear behaviors (3–7). Briefly, fear conditioning focuses on how an initially innocuous conditioned stimulus (CS; e.g., auditory, visual, contextual cues), upon pairing with a noxious unconditioned stimulus (US; usually electric shock) that reflexively elicits unconditioned response (UR; namely defensive reactions), becomes capable of eliciting conditioned response (CR; e.g., freezing in rodents, increased skin conductance in humans). A century of fear conditioning research has led to wide-ranging discoveries. In particular, fear conditioning experiments have fundamentally transformed learning theories from the archaic contiguity (or temporal) relationship (8–10) to the modern contingency (or informational) relationship between the CS and US (11–14), revealed detailed neurobiological mechanisms of learning and memory (15–17) and influenced contemporary cognitive behavioral therapy for various anxiety and traumatic-stressor related disorders, such as panic, phobic and posttraumatic stress disorders (18–22).

Despite its utility and appeal, fear conditioning paradigms nonetheless simplify behavioral analyses of fear, ignoring the multitude of actions and decisions that animals and humans utilize to survive the breadth of risky situations in the real world (23–28). Moreover, the prevalent notion that fear conditioning produces adaptive associative fear memory has yet to be ecologically validated. In fact, some researchers have questioned the evolutionary logic underlying fear conditioning; “No owl hoots or whistles 5 seconds before pouncing on a mouse…Nor will the owl give the mouse enough trials for the necessary learning to occur…What keeps animals alive in the wild is that they have very effective innate defensive reactions which occur when they encounter any kind of new or sudden stimulus” (29). Indeed, laboratory rodents exhibit unlearned, instinctive fear responses to advancing artificial terrestrial and aerial predators (30, 31), overhead looming stimuli (32), and predator odors (33).

Here, we investigated for the first time whether fear conditioning readily transpires and modifies subsequent behavior of animals in a naturalistic environment. To achieve this, hunger-motivated rats searching for a food pellet in a large arena—a purposive behavior (34)—were presented with a discrete tone CS followed by a painful US to their dorsal neck/body region by means of chronically implanted subcutaneous wires (Fig. 1A). A dorsal neck/body shock better simulates real predatory strike compared to footshock used in standard fear conditioning studies, as it is unlikely that predators direct their attacks on small prey animal’s paws. Additionally, in nature, bodily injuries are normally inflicted by external agents (namely, predators in animals and perpetrators in humans). Thus, other groups of rats were presented with a looming aerial predator (i.e., a lifelike great horned owl) preceded with and without a tone CS and followed by the same US (Fig. 1B-D). A single trial tone-shock, tone-owl, tone/owl-shock and owl-shock training was employed because multiple bodily harm encounters would prove fatal in nature, antithetical to the natural selection of fear conditioning (29). Later, all animals’ reactions to the tone cue were examined while foraging for food in the open arena.

**Fig. 1.**
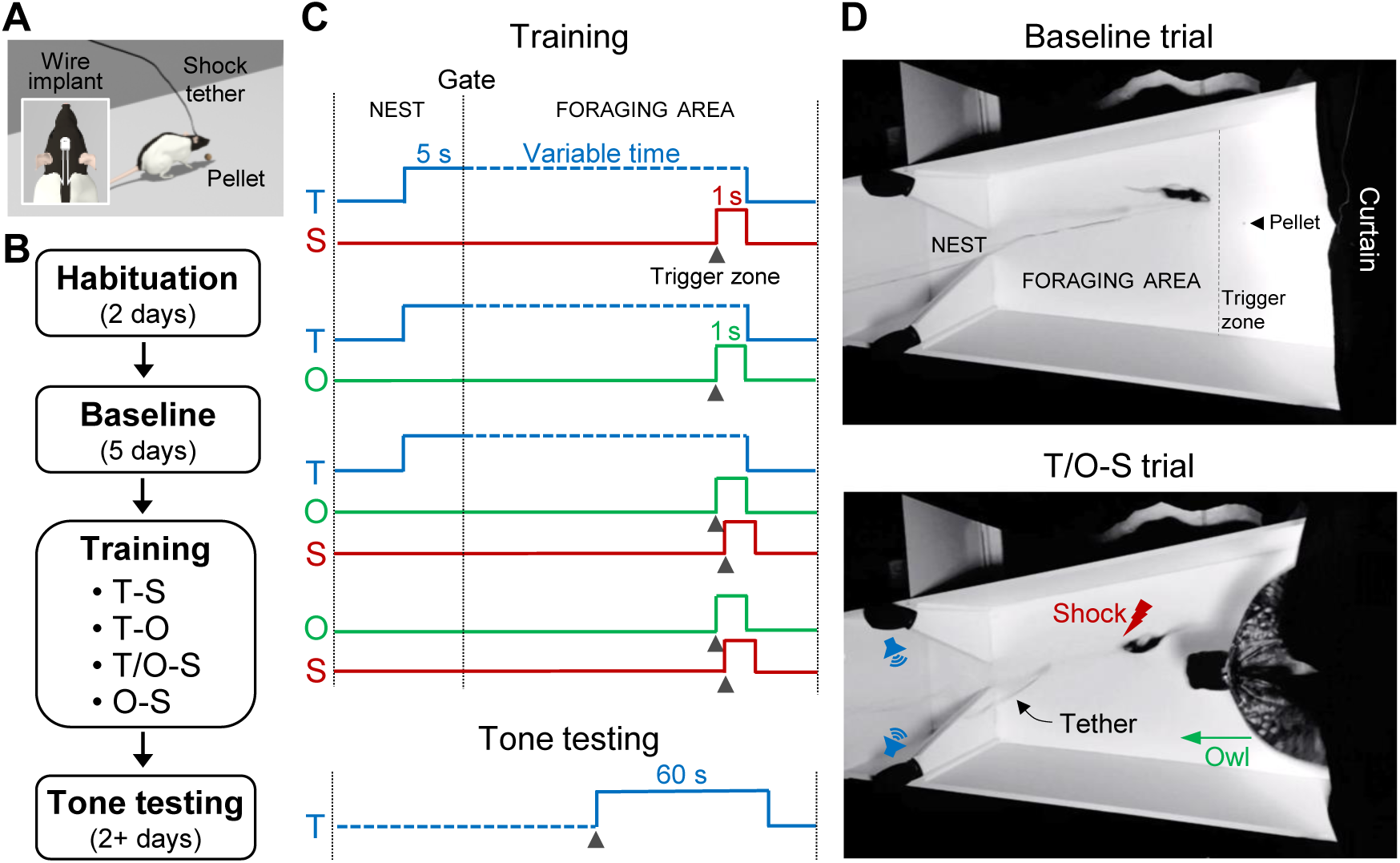
Experimental design of fear conditioning in a naturalistic setting. **(A)** An illustration of a tethered rat foraging for a food pellet in the open arena (inset shows a headstage and placement of subcutaneous shock wires). **(B)** Timeline of experiment. *Habituation*: Rats were placed in a closed nest with dispersed food pellets for 30 min/day. *Baseline*: Rats were allowed to leave the nest to discover food pellets placed 25-125 cm (in 25 cm increments from the nest) in the foraging arena. *Training*: Animals approaching the pellet location experienced a delayed pairing of tone-shock (T-S), tone-owl (T-O), tone/owl-shock (T/O-S), or owl-shock (O-S). *Tone Test*: On subsequent days, all rats were placed back in the foraging arena and upon nearing the food pellet, the tone was activated. **(C)** Schemas of delayed pairings of stimuli. The T-S, T-O and T/O-S (but not O-S) groups were presented with a tone 5 s before the gate opening that stayed on until the animals were within 25 cm of the food pellet, at which the tone co-terminated with the triggered shock (1 s), owl (1 s) or owl-shock (100 ms interstimulus interval, ISI) stimuli. **(D)** A representative rat in the foraging arena (208 cm length x 66-120 cm expanding width x 61 cm height) during a baseline trial, where the animal successfully acquires the pellet, and during a T/O-S trial, where the animal flees from looming owl and shock into the nest (69 cm length x 58-66 cm width x 61 cm height).

## Results

### Baseline foraging

Female and male rats were pseudo-randomly assigned to tone-shock (8 females, 8 males), owl-shock (8 females, 8 males), tone/owl-shock (6 females, 8 males), and tone-owl (4 females, 4 males) groups and implanted with subcutaneous wires in their dorsal neck/body (Fig. 1A-C). After recovery from the surgery, the rats were trained to exit a nest compartment upon gate opening to procure a sizable 0.5 g food pellet placed at variable distances in a large, expanding open arena (Fig. 1D, top panel). On the first baseline day, female rats took a significantly longer amount of time to procure the food pellet compared to male rats (*Supplementary materials*, Fig S1, Baseline day 1). This initial difference in foraging behavior likely represents heightened spatial neophobia (risk-averse to novel environments) in female rats. As rats became familiar with the foraging arena, the latency and duration measures declined across 5 baseline days comparably in both sexes, with no further statistical differences in latencies for pellet procurement. Because there were no reliable sex differences in subsequent fear conditioning dependent variables (*Supplementary materials*, Fig. S2), the four groups were collapsed across sexes.

### Fear conditioning

On the training day, all rats first underwent three foraging trials with pellets fixed at the longest distance (125 cm) to confirm comparable pre-fear conditioning foraging behavior between groups (Fig. 2A, Baseline). Afterwards, animals were exposed to a tone-shock, an owl-shock, a tone/owl-shock or a tone-owl pairing in the manner shown in Fig. 1 (*Supplementary materials*, Movie S1). Those rats presented with the tone CS 5-sec prior to the gate opening (i.e., tone-shock, tone-owl, tone/owl-shock groups) took more time to enter the foraging arena in comparisons to owl-shock animals unexposed to the tone (Fig. 2B, Leave nest latency); this indicates that the tone was a salient cue that animals were attentive to and thus conditionable. Once in the foraging arena, all animals readily advanced toward the pellet and breached the trigger zone (25 cm from the pellet) to activate the shock, owl, or owl-shock stimuli (Fig. 2B, Trigger zone latency). In response to the shock, owl, or owl-shock, all rats promptly fled from the foraging arena to the nest (Fig. 2B, Escape latency; Fig. 2D,E, Escape speed). Figure 2C shows representative track plot examples of tone-shock, owl-shock, tone/owl-shock and tone-owl animals successfully procuring the pellet during pre-tone baseline but not during tone conditioning. The fact that the escape latency and running speed were not significantly different between the tone-owl and other groups indicates that the looming owl-induced innate fear sans pain was just as effective in eliciting the flight UR as the painful shock or shock-owl combination. However, inspections of the escape trajectories revealed that the tone-shock and tone-owl groups tended to flee linearly to the nest, whereas the owl-shock and tone/owl-shock groups that experienced a dorsal neck/body shock 100 ms after the looming owl (mimicking realistic predatory attack) and begun their flight to the nest inclined to escape circuitously (Fig. 2F,H). This was supported by significant group differences in the escape distances (Fig. 2G) and trajectory angles (Fig. 2I), where owl-shock and tone/owl-shock groups traveled longer distances and had higher angle variances, respectively, during their escape routes than tone-shock and tone-owl groups.

**Fig. 2.**
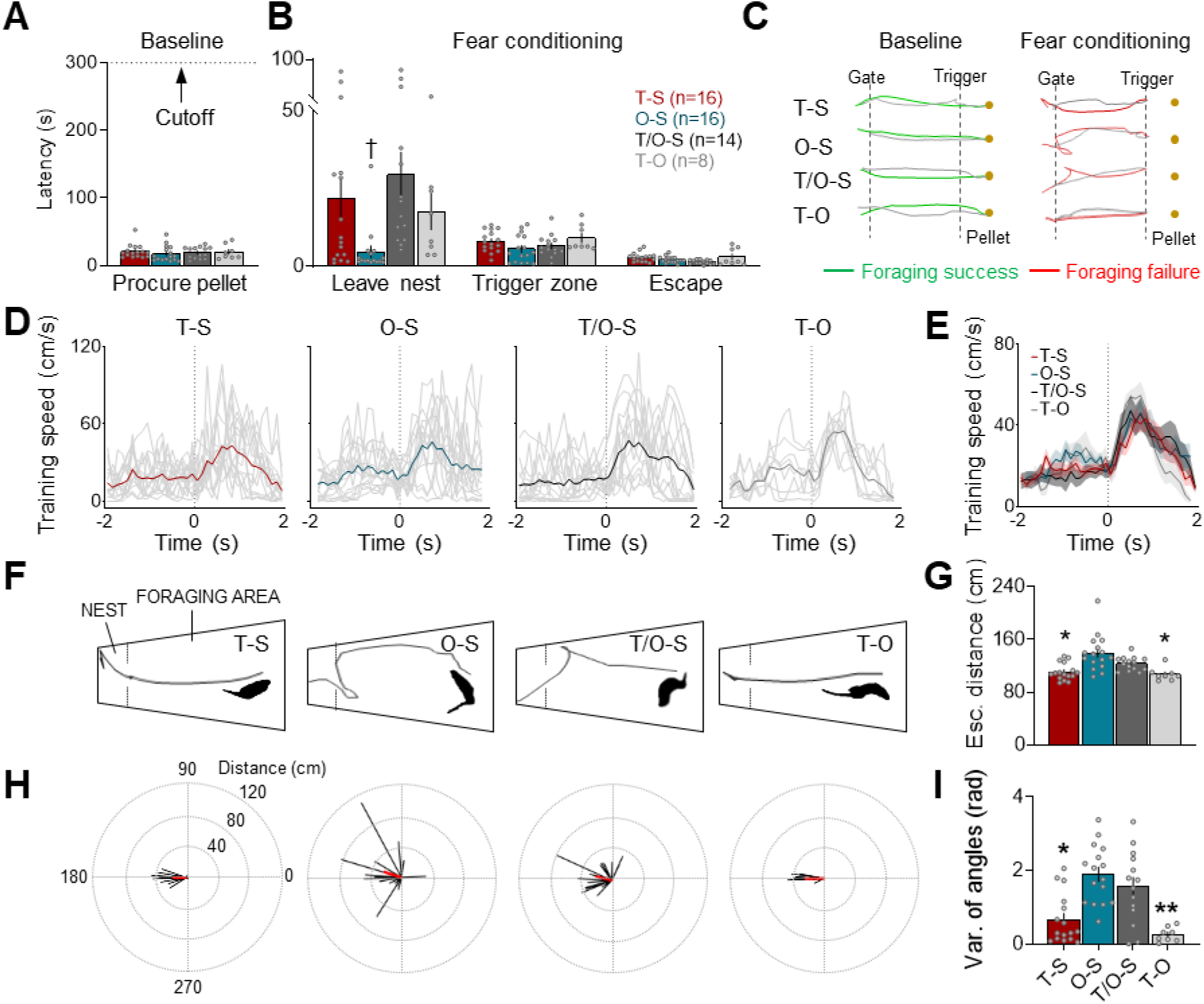
Foraging and escape behaviors during fear conditioning. **(A)** Pre-conditioning baseline latencies (mean ± SEM) to procure food pellets in the foraging arena were equivalent between T-S (red), O-S (blue), T/O-S (dark gray) and T-O (light gray) groups (Kruskal-Wallis, H = 2.694, *p* = 0.441). **(B)** During fear conditioning, the T-S, T/O-S and T-O groups exposed to the tone 5 s before the gate opening had significantly longer latencies to leave the nest than the O-S group (left panel, Kruskal-Wallis, H = 18.6, *p* < 0.001; pairwise comparisons, *p* = 0.008 for T-S vs. O-S, *p* = 0.011 for O-S vs. T-O, *p* < 0.001 for O-S vs. T/O-S, *p* = 0.69 for T-S vs. T-O, *p* = 0.631 for T-S vs. T/O-S, *p* = 0.343 for T/O-S vs. T-O). Once outside the nest, however, the latency to breach the trigger zone, enroute to the pellet, was not reliably different among the groups (Kruskal-Wallis, H = 7.453, *p* = 0.059). In response to the triggered shock, owl or owl-shock, all groups showed similar escape-to-nest latencies (Kruskal-Wallis, H = 6.141, *p* = 0.105). **(C)** Representative track plot examples from T-S, O-S, T/O-S and T-O animals during the baseline, when animals successfully procured the pellet, and during the fear conditioning, when the same animals fled from shock, owl or owl-shock stimuli and thus unable to attain the pellet. **(D)** Mean instantaneous speed (± SEM) of each group 2 sec before and after the shock, owl or owl-shock onset (t = 0). Thin, grey lines represent individual animal data. **(E)** All groups showed comparable escape speed to the shock, owl, and owl-shock stimuli (Kruskal-Wallis, H = 0.901, *p* = 0.825). **(F)** Representative track plots showing escape paths of T-S, O-S, T/O-S and T-O animals. The inset silhouette images show that the T-S and T-O animals were facing forward at the time of the shock or owl stimulus whereas the O-S and T/O-S animals were turning back at the time of the shock stimulus because of the 100 ms owl-shock interstimulus interval. **(G)** Mean escape distance (± SEM) from the trigger zone to the nest. The O-S and T/O-S groups travelled longer distances to escape compared to the T-S and T-O groups (Kruskal-Wallis, H = 21.98, *p* < 0.001; pairwise comparisons, *p* = 0.014 for T-S vs. T/O-S, *p* = 0.008 for T/O-S vs T-O, *p* = 0.001 for T-S vs. O-S, *p* = 0.001 for O-S vs T-O). **(H)** Representative vector plots of each group showing variabilities in their escape paths. **(I)** Mean variance (± SEM) of escape trajectory angles (radian) from the trigger zone to the nest. The O-S and T/O-S groups had greater variance in their escape trajectories when fleeing back to the nest (Kruskal-Wallis, H = 22.37, *p* < 0.001; pairwise comparisons, *p* = 0.022 for T-S vs. T/O-S, *p* = 0.003 for T/O-S vs T-O, *p* = 0.002 for T-S vs. O-S, *p* < 0.001 for O-S vs T-O). († compared to T-S, T/O-S, and T-O; * compared to O-S and T/O-S, *p <* 0.05, ** *p <* 0.01, *** *p* < 0.001; # compared to T/O-S, *p* < 0.05, *## p <* 0.01).

### Context (pre-tone) testing

On the following day, animals were placed back in the nest and underwent three pre-tone baseline trials (maximum 300 sec to retrieve the food pellet placed at 125 cm) to assess whether previous encounters with tone-shock, owl-shock, tone/owl-shock and tone-owl stimuli combinations produced fear of the arena. As can be seen in Figure 3A, the owl-shock and tone/owl-shock groups took significantly longer latencies to procure the pellet (i.e., the time from gate opening-to-return to nest with the pellet) than the tone-shock and tone-owl groups on the first day of testing. The lengthened times to enter the foraging arena exhibited by owl-shock and tone/owl-shock rats likely reflect inhibitory avoidance resulting from the previous predatory attack experience in the arena (35). In contrast, the fact that the pre-tone test baseline latencies of tone-shock and tone-owl rats (*Supplementary materials*, Fig. S3) were not reliably different from their baseline latencies from the fear conditioning day (prior to experiencing tone-shock or tone-owl) suggests that contextual fear conditioning failed to transpire in these animals despite their robust escape behavior to tone-shock and tone-owl experiences. Similar patterns of group differences, albeit lesser magnitudes, were observed on the second day of pre-tone baseline trials (Fig. 3C).

**Fig. 3.**
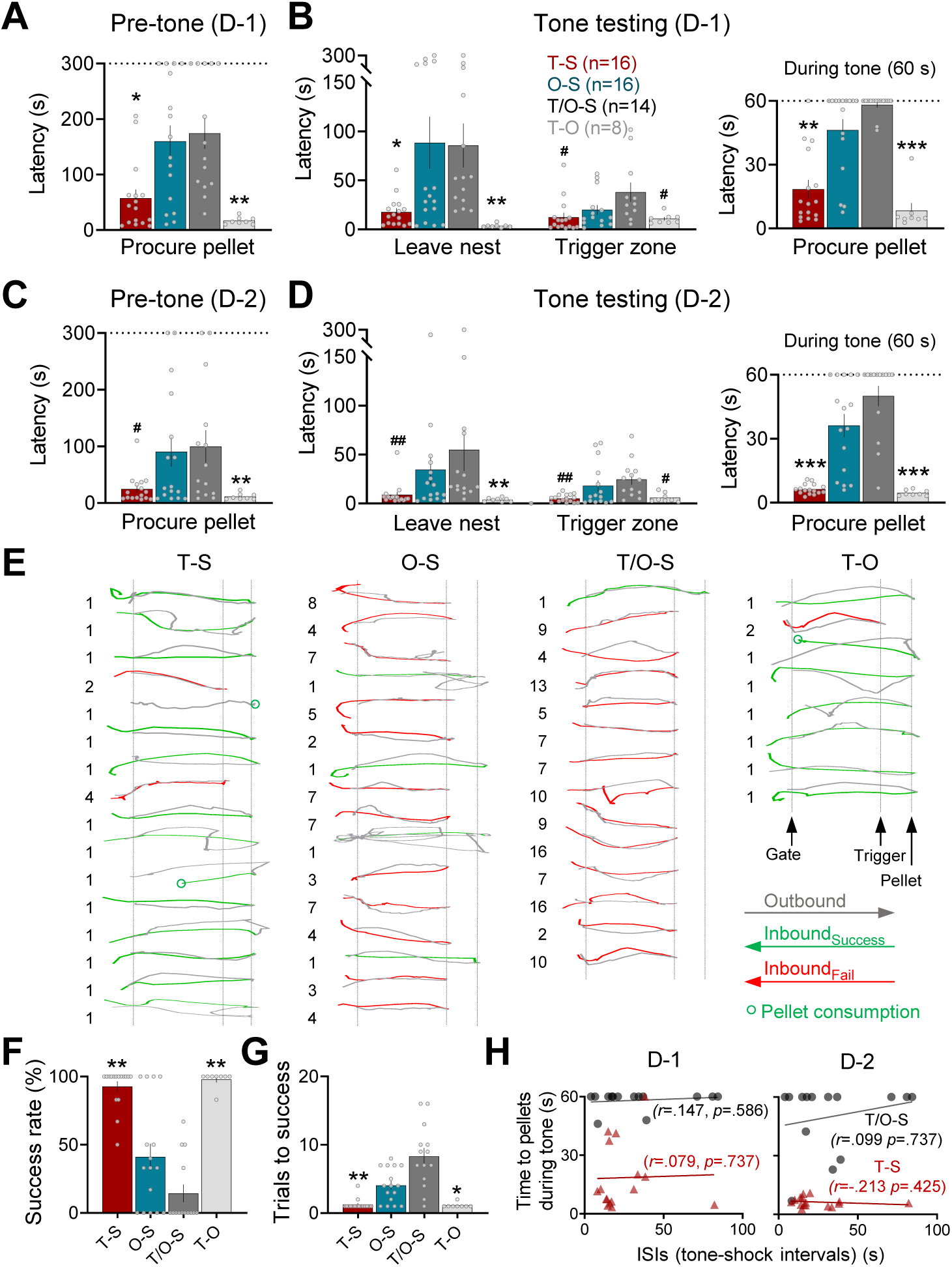
Foraging and escape behaviors during tone testing. **(A)** The mean latency (± SEM) to procure the pellet during the pre-tone baseline trials on testing day 1 (D-1). Both O-S and T/O-S groups took significantly longer times to exit (gate opening, t=0) and return to the nest with the pellet than T-S and T-O groups (Kruskal-Wallis, H = 20.518, *p* < 0.001; pairwise comparisons, *P* = 0.003 for T-S vs. T/O-S, *p* < 0.01 for T/O-S vs. T-O, *p* = 0.013 for T-S vs. O-S, *p* < 0.001 for O-S vs. T-O). **(B)** The times (mean ± SEM) to leave nest and reach trigger zone on day 1 tone test trials. Both O-S and T/O-S groups had longer latencies to leave nest (Kruskal-Wallis, H = 27.071, *p* < 0.001; pairwise comparisons, *p* = 0.003 for T-S vs. T/O-S, *p* < 0.001 for T/O-S vs. T-O, *p* = 0.044 for T-S vs. O-S, *p* < 0.001 for O-S vs. T-O. Once outside the nest, the T/O-S group took longer time to reach the trigger zone than the T-S and T-O (Kruskal-Wallis, H = 9.153, *p* = 0.027; pairwise comparisons, *p* = 0.019 for T-S vs. T/O-S, *p* = 0.042 for T/O-S vs. T-O). During the tone test, the latencies to procure the pellet within the 60 s allotted time were significantly longer in O-S and T/O-S animals compared to T-S and T-O animals (Kruskal-Wallis, H = 34.428, *p* < 0.001; pairwise comparisons, *p* < 0.001 for T-S vs. T/O-S, *p* < 0.001 for T/O-S vs. T-O, *p* = 0.02 for T-S vs. O-S, *p* < 0.001 for O-S vs. T-O). **(C)** The mean latency (± SEM) to procure the pellet during the pre-tone baseline trials on testing day 2 (D-2). O-S and T/O-S groups continued to have longer latencies to exit (gate opening, t=0) and return to the nest with the pellet than T-S and T-O groups (Kruskal-Wallis, H = 12.47, *p* = 0.006; pairwise comparisons, *p* = 0.022 for T-S vs. T/O-S, *p* = 0.002 for T/O-S vs. T-O, *P* = 0.009 for O-S vs. T-O). **(D)** The times (mean ± SEM) to leave nest and reach trigger zone on day 2 tone test trials. There were group differences in the latencies to leave nest (Kruskal-Wallis, H = 21.505, *p* < 0.001; pairwise comparisons, *p* = 0.001 for T-S vs. T/O-S, *p* < 0.001 for T/O-S vs. T-O, *p* = 0.002 for O-S vs. T-O). Once outside the nest, there were group differences in the latencies to reach the trigger zone (Kruskal-Wallis, H = 21.531, *p* < 0.001; pairwise comparisons, *p* < 0.001 for T-S vs. T/O-S, *p* < 0.001 for T/O-S vs. T-O, *p* = 0.037 for O-S vs. T-O). During the tone test, the latencies to procure the pellet within the 60 s allotted time were significantly longer in O-S and T/O-S animals compared to T-S and T-O animals (Kruskal-Wallis, H = 37.223, *p* < 0.001; pairwise comparisons, *p* < 0.001 for T-S vs. T/O-S, *p* < 0.001 for T/O-S vs. T-O, *p* < 0.001 for T-S vs. O-S, *p* < 0.001 for O-S vs. T-O). **(E)** Individual track plots during the first tone exposure from all animals from each group. The parenthesized numbers next to plots represent the trial(s) needed for successful foraging. **(F)** The overall success rates of procuring the pellet on the first testing day were significantly lower in the O-S and T/O-S groups compared to the T-S and T-O groups (Kruskal-Wallis, H = 32.299, *p* < 0.001; pairwise comparisons, *p* < 0.001 for T-S vs. T/O-S, *p* < 0.001 for T/O-S vs. T-O, *p* = 0.001 for T-S vs. O-S, *p* = 0.003 for O-S vs. T-O). **(G)** The O-S and T/O-S animals required extended trials to obtain the pellet (Kruskal-Wallis, H = 32.004, *p* < 0.001; pairwise comparisons, *p* < 0.001 for T-S vs. T/O-S, *p* < 0.001 for T/O-S vs. T-O, *p* = 0.002 for T-S vs. O-S, *p* = 0.011 for O-S vs. T-O). **(H)** In T-S and T/O-S animals, there were no reliable correlations (Spearman’s correlation coefficient) between the tone-induced suppression of pellet procurement (an index of fear) and the temporal intervals (i.e., ISIs) between tone CS onset and shock US onset in neither testing day 1 nor 2. (* compared to both O-S and T/O-S, *p <* 0.05, ** *p <* 0.01, *** *p* < 0.001; # compared to T/O-S, *p* < 0.05, *p <* 0.01).

### Tone testing

Immediately after the pre-tone baseline, all groups were subjected to three successive tone test trials (one minute apart). The owl-shock and tone/owl-shock animals continued to take longer latencies to exit the nest compared to tone-shock and tone-owl animals (Fig. 3B, Leave nest latency). Once in the foraging arena, the tone/owl-shock group’s latency to approach 25 cm from the pellet to trigger the tone were marginally but reliably longer than those of tone-shock and tone-owl groups, but not owl-shock group (Fig. 3B, Trigger zone latency). Upon the activation of tone (60 s continuous), the majority of owl-shock and tone/owl-shock animals promptly fled to the nest, thereby significantly increasing the latency to procure the pellet (60 s = unsuccessful), whereas the tone-shock and tone-owl animals were largely unaffected by the tone and readily procured the pellet (Fig. 3B, Procure pellet latency). The second day of tone testing yielded similar patterns of group differences (Fig. 3D). Figure 3E shows individual track plots from all animals with the initial number of trial(s) necessitated for successful foraging. Further analyses across tone testing days (3 trials/day) showed that the overall success rates of procuring the pellet were significantly lower in owl-shock and tone/owl-shock groups compared to tone-shock and tone-owl groups (Fig. 3F), and that owl-shock and tone/owl-shock animals required extended trials to reliably obtain the pellet (Fig. 3G). Because the temporal interval between the CS and US is well known to be crucial in various types of Pavlovian conditioning, including fear conditioning (36), we examined whether tone fear conditioning transpired in a specific (optimal) range of interstimulus intervals (ISI) but was masked by non-optimal ISIs. We found no significant correlation between the ISIs and the magnitudes of tone-induced suppression of pellet procurement in tone-shock animals, indicating that tone fear conditioning failed to materialize across varying ISIs of delay conditioning (Fig. 3H). Conversely, in the tone/owl-shock animals, the tone-induced suppression of pellet procurement was uniformly observed across different ISIs, suggesting that the observed fear in these animals may not necessarily reflect Pavlovian conditioning (Fig 3H). These results of delayed tone-shock paired animals failing to show conditioned tone fear and contextual fear suggest that standard fear conditioning does not readily occur in naturalistic environment. Instead, the finding of owl-shock animals displaying robust fear to a novel tone, which the animals never heard before, suggests that non-associative sensitization-like processes play a crucial role in protecting animals in the real world.

## Discussion

It is generally believed (though never validated) that there is behavioral continuity of Pavlovian fear conditioning from the laboratory to real-life situations, and thus understanding the mechanisms of fear conditioning will have clinical relevance. The present study directly investigated whether fear conditioning readily occurs in naturalistic situations that animals are likely to encounter in their habitats. Standard fear conditioning in rodents takes place in small experimental chambers, and several studies have shown that a single tone CS-footshock US pairing (i.e., delay fear conditioning) reliably produces conditioned freezing in rats and conditioned tachycardia/freezing in mice (37). One-trial delay tone fear conditioning has also been demonstrated in human subjects using a loud white noise US and assessing conditioned skin conductance response (38). However, in the present study, where rats are exhibiting a purposive foraging behavior (34) in a large arena, a delayed pairing of tone CS and dorsal neck/body shock US (tone-shock group) produced virtually no evidence of auditory (and contextual) fear conditioning across a range of CS durations (i.e., ISIs). A similar pairing of tone CS and looming owl (tone-owl group) also failed to produce auditory fear conditioning despite the owl US evoking robust fleeing UR. In contrast, foraging rats that experienced a looming owl and shock pairing (owl-shock group) later exhibited robust fear (escape) behavior to a novel tone presentation. In the tone/owl-shock animals, the escape behaviour was uniformly observed across different ISIs, suggesting that the observed fear to the tone stimulus in this group may not be a Pavlovian response. These findings then point to a nonassociative sensitization (or sensitization-like) process, rather than associative fear conditioning, as playing a vital function in risky (i.e., predatory attack) situations that animals encounter in nature.

The tone CS (3 kHz, 80 dB, ranging 9-86.6 s) and shock US (2.5 mA, 1 s) employed in the present study were effective in eliciting orienting and fleeing responses, respectively, and were presented to animals in the manner (i.e., a delay conditioning) that satisfied the stimuli saliency, intensity, surprising, and temporal contiguity requirements for conditioning (39–41). Then, what can account for one-trial auditory fear conditioning, demonstrated in standard Pavlovian paradigms (35, 37, 38, 42), not emerging in animals that left the safe nest to forage for food in an open arena? It may well be that rats are not biologically predisposed to associate discrete CS and US in natural (complex) environments where competing hunger-driven and fear-driven motivated behaviors are freely expressed. Indeed, in real-life, only a small minority of people experiencing trauma develop posttraumatic stress disorder (PTSD) and even with re-exposure to the same trauma there is low incidence PTSD (43, 44). In contrast, standard experimental chambers may be conducive to fear conditioning because they are simple and limit the repertoire of behaviors. The absence of one-trial fear conditioning in a naturalistic setting may be analogous to “The Rat Park Experiment,” where rats housed in an enriched environment with plants, trees and social interaction resist drug addiction behavior evident in standard cage-housed rats (45, 46). Animals tested in naturalistic paradigms are given choices that do not force their behaviors into dichotomies (i.e., freezing or no freezing; drug craving or no drug craving). Allowing for an expanded behavioral repertoire, while more difficult to study, may thus yield a greater understanding of behaviors and their underlying brain mechanisms.

It should also be noted that fear encounters in real life generally occur in the presence of external agents or forms (i.e., predators/conspecifics in animals and assailants/combatants in humans), which is virtually nonexistent in standard Pavlovian fear conditioning paradigms. Thus, the effects of a discernable entity in associative fear learning have never been investigated. By simulating a realistic life-threatening situation, i.e., a looming aerial predator that instinctively elicited flight behavior followed by somatic pain, we found that rats engaged in purposive behavior utilize nonassociative sensitization as their primary defensive mechanism. The fact that the owl-shock and tone/owl-shock animals exhibited relatively nonlinear, erratic escape trajectories to the nest compared to linear escape trajectories in tone-shock animals (Fig. 2F-I) suggests the intriguing possibility that the same dorsal neck/body shock US may be interpreted as a life-or-death (panic) situation in the presence of an external threat agent versus a mere startling (nociceptive) situation in the absence of an external threat agent. The erratic flight behavior in the presence of a looming owl may represent the penultimate stage of circa-strike, or “life-or-death,” behavior within the “predatory imminence continuum” theory (47). Functionally, a sensitized fear system may intensify avoidance behavior, which in turn effectively transposes novel, neutral cues into “false positives” to prioritize survival in natural environment (29). In other words, nonspecific sensitization-based overestimation of danger may be a more prudent course for survival than relatively more specific association-based prediction of danger.

Clark Hull (48) has posited that Pavlovian fear conditioning offers biological utility by circumventing a “bad biological economy” of defense reaction always necessitating injury. This prevailing view that ascribes preeminent importance of fear conditioning as the primary defensive mechanism is likely to be a theoretical simplification and provides an incomplete picture of fear, as its function in a natural environment may be rather limited (i.e., lacks face validity). It may well be possible to produce fear conditioning in naturalistic settings with further CS-US trials but then this too would be a bad biological economy as such learning will dramatically reduce biological fitness. It is also important to recognize inconsistencies in the literatures, such as clinical studies that have reported that patients with anxiety disorders, such as phobias, have trouble recalling the particular pairing of the fear event with its aversive consequences (49, 50). The increased utilization of naturalistic fear paradigms that simulate dangers that animals and humans encounter in real life will enable us to clarify, update, and revise fear concepts derived largely from fear conditioning studies and in doing so facilitate future progress in the treatment of fear disorders.

## Materials and Methods

### Subjects

Sixty-two Long-Evans rats (3-4 months old; 32 females and 30 males), purchased from Charles-Rivers Laboratories, were initially pair-housed by sex for 5-7 days of acclimatization in a climate-controlled vivarium (accredited by the Association for Assessment and Accreditation of Laboratory Animal Care), with a reversed 12-h light/dark cycle (lights on at 7 PM). After undergoing subcutaneous wire implant surgery (described below), animals were individually housed and placed on a standard food-deprivation schedule with *ad lib* access to water to gradually reach and maintain ∼85% normal body weight. All experiments were performed during the dark phase of the cycle in strict compliance with the University of Washington Institutional Animal Care and Use Committee guidelines.

### Surgery

Under isoflurane anesthesia, rats were mounted on a stereotaxic instrument (Kopf), and two Teflon-coated stainless-steel wires (0.0003 inch bare, 0.0045 inch coated; A-M Systems, Everett, WA) were inserted in the dorsal neck/back region of body. The wire tips were exposed (∼1 cm), bent to a V-shape, and hooked to subcutaneous tissue (36). The other ends of the wires were affixed to a headstage (Plastics One, MS303-120), which was then cemented to the animal’s skull embedded with 6 anchoring screws. While still under anesthesia, animals were connected to a shock-apparatus and given a mild shock to observe muscle twitching; 6 rats that showed no reaction to shock were removed from the experiment. Animals were given 4 days of postoperative recovery and were adapted to handling for 5 days before nest habituation.

### Foraging Apparatus and Stimuli

A custom-built foraging arena consisted of a nest (69 cm length x 58-66 cm width x 61 cm height) that opened via an automated sliding gate to reveal a large, expanded foraging area (208 cm length x 66-120 cm width x 61 cm height) where 0.5 g food pellets (grain-based; F0171, Bio-Serv) were placed at variable locations (Fig. 1A). The testing room was kept under red light (11 lux foraging area, 2 lux nest area) with constant white noise (72 dB) playing in the background. Prior to placing each animal, the arena was wiped with 70% ethanol. The ANY-maze software and Ami interface system (Stoelting) connected to a PC automatically tracked the animal’s position in the arena, via a ceiling mounted camera, and triggered the tone, shock and aerial predator stimuli: (i) 3 kHz, 80 dB tone CS was produced using Anymaze (Stoelting) and presented through two speakers mounted on the nest-foraging border; (ii) 1 s, 2.5 mA shock US was delivered to the animal’s dorsal neck/back region via a headstage tethered to a stimulus-isolator (Bak); (iii) A life-like model owl (31), mounted onto a 92 cm pneumatic air cylinder (Bimba) at the opposite end of the foraging arena and hidden behind a black curtain, plunged downward towards the rat (46 cm/s), then retracted back to it starting position.

### Behavioral Procedure

Upon reaching and maintaining 85% normal body weight, animals were transported to the experimental room and underwent series of habituation, baseline, fear conditioning, and testing sessions.

(*Habituation days*) Animals were placed in the nest scattered with 20 food pellets (0.5 g, grain-based, Bio-Serv) for 30 min/day for 2 consecutive days to acclimatize and associate the nest with food consumption.

(*Baseline days*) After 1 minute in the nest sans food pellets, the gate opened, and the animal was allowed to explore the large foraging arena and find a pellet placed 25 cm away from the nest (first trial). As soon as the animal took the sizeable 0.5 g pellet back to the nest, the gate closed. Once the animal finished eating, the second trial with the pellet placed 50 cm and then the third trial with the pellet placed 75 cm commenced in the same manner. Animals underwent 3-5 consecutive baseline days, with the pellet distances gradually extending to 75, 100 and 125 cm, and they were also accustomed to tethering beginning on baseline day 3 onward.

(*Fear conditioning day*) Rats, pseudo-randomly assigned into tone-shock, tone-owl, tone/owl-shock and owl-shock groups (Fig. 1), underwent 3 baseline trials with the pellet placed at 125 cm from the nest. On the 4^th^ trial, the tone-shock, tone-owl and tone/owl-shock animals were exposed to a tone CS that came on 5 seconds before the gate opened and remained on until they reached the trigger zone (25 cm to the pellet). For tone-shock and tone-owl animals, the tone co-terminated with the shock US and the owl looming, respectively. For tone/owl-shock animals, the shock occurred 0.1s sec after the owl looming and co-terminated with the tone. Two animals in the tone/owl-shock group were excluded because they failed to leave the nest within 2 min. The owl-shock animals were subjected to the same owl looming-shock pairing (as the tone/owl-shock animals) but in the absence of tone. All rats fled to the nest in reaction to the shock and/or looming owl, at which time the gate was closed. After 1 minute in the nest, the animals were placed back into their homecage.

(*Testing days*) All rats underwent 3 baseline trials (a maximum of 300 sec to retrieve the pellet) to assess whether shock and/or looming owl encounter the previous day resulted in the fear of the arena (i.e., contextual fear). Afterwards, animals were presented with the tone cue when they approached the trigger zone (25 cm to the pellet). The tone played continuously for 60 sec, after which the tone test trial ended. Animals underwent 3 tone tests daily until they successfully attained the pellet (i.e., fear extinction).

### Data Analyses

Statistical analyses were performed using SPSS (IBM, version 19) and R (The R Foundation, version 3.5.3). Body tracking positions were obtained using Deep Lab Cut (51) and analyzed using a self-written script in Python (Python Software Foundation). Animal sample sizes were determined using a power analysis performed by G*Power (G*Power, version 3.0.1, Franz Faul; power=0.95, alpha=0.05, effect size=0.5, two-tailed). A Levene’s test for normality showed significance for the data, thus nonparametric tests were used for analysis. Because there were no significant sex differences in any stages of the experiment after the first day of baseline (*Supplementary materials*, Fig. S1), data from females and males were pooled together for all analyses (*Supplementary materials*, Fig. S2). Statistical significance was set at *P* < 0.05. Graphs were made using GraphPad Prism (version 8).

### Data Availability

The data that support the findings of this study and the relevant analysis code are available from the Dryad data repository. https://doi.org/10.5061/dryad.76hdr7sxk Reviewer Link: https://datadryad.org/stash/share/0O_D25HmXortJJoB9bMz5YMUvKOM09RLtEv-TOR2sRc

## Acknowledgments

We thank Lori A. Zoellner for valuable comments on the manuscript, and Heather Wu for assistance in the experiment. This study was supported by National Institutes of Health Grant MH099073 (to J.J.K.).

## Supplementary Information for

**Fig. S1.**
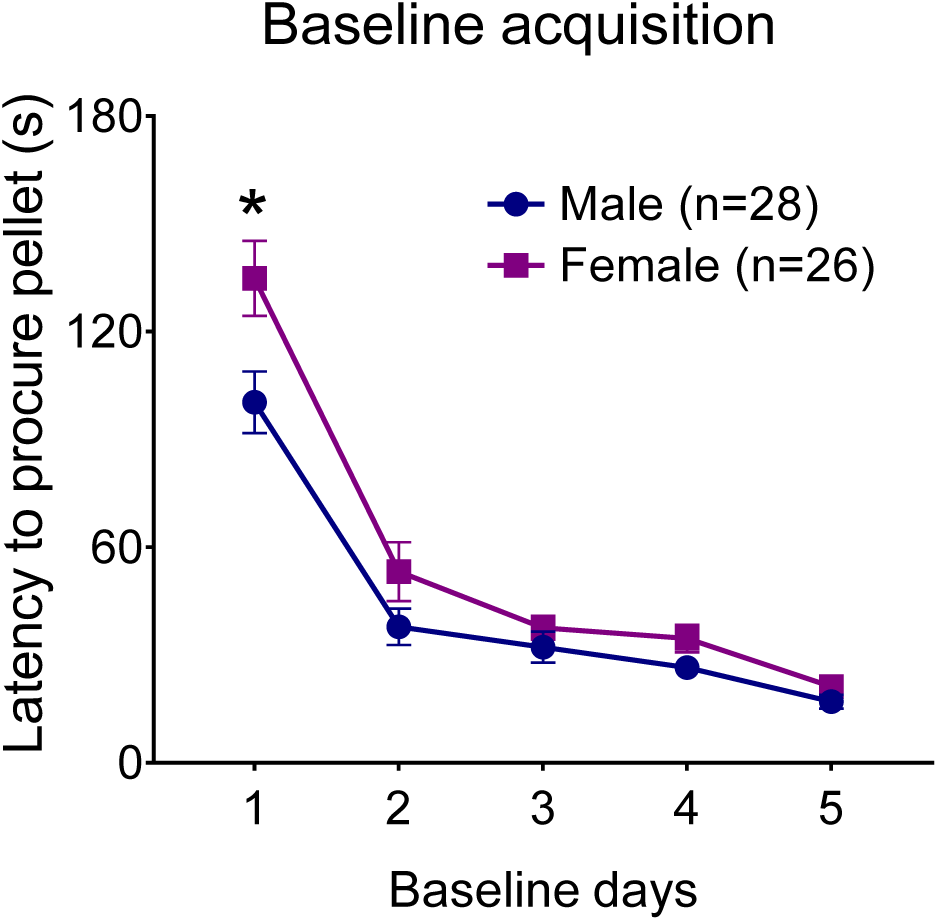
Initial sex differences in the baseline latency to procure pellets. Average latencies (±SEM) to procure food pellets in foraging area. Females had longer latencies to procure pellets than males during the first baseline session day 1 (Mann-Whitney U, z = 2.476, *p* = 0.013) but not subsequent baseline session days 2-5 (Mann-Whitney U, Baseline 2: z = 1.039, *p* = 0.299; Baseline 3: z = 1.922, *p* = 0.055; Baseline 4: z = 1.112, *p* = 0.266; Baseline 5: z = 1.904, *p* = 0.057). *\* p* < 0.05.

**Fig. S2.**
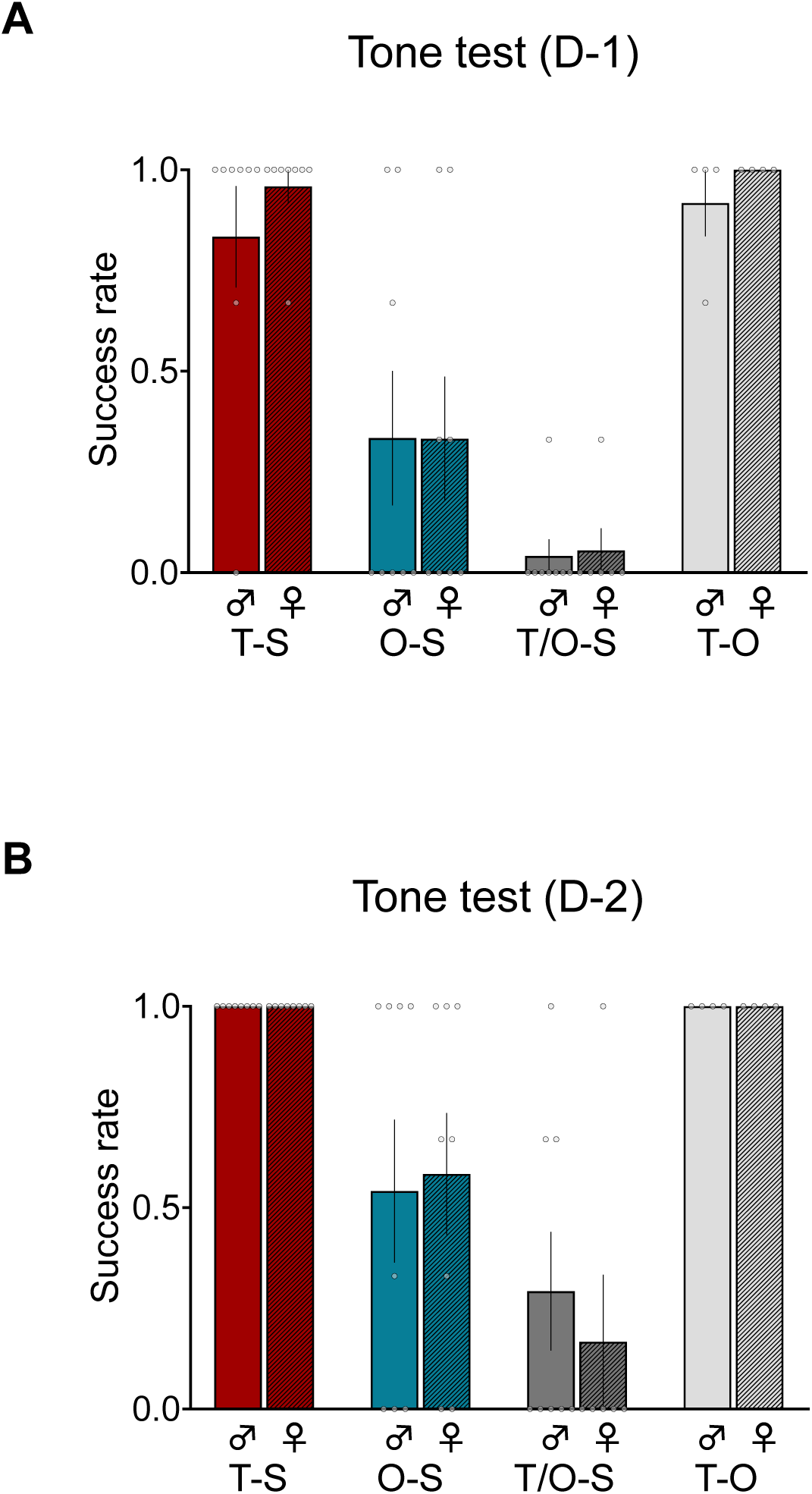
No reliable sex differences in the procurement of pellets during tone tests. **(A)** Mean (± SEM) success rate for procuring food pellets during the first day of tone testing. No significant differences were found between males and females in tone-shock (T-S), owl-shock (O-S), tone/owl-shock (T/O-S) and tone-owl (T-O) groups (Mann-Whitney U; z = 0.694, *p* = 0.645 for T-S; z = 1.0, *p* = 0.317 for O-S; z = 0.212, *p* = 1.0 for T/O-S; z = 0.234, *p* = 0.815 for T-O). **(B)** Mean (± SEM) success rate for procuring food pellets during the second day of tone testing. No sex differences were observed in all groups (Mann-Whitney; z = 0, *p* = 1.0 for T-S; z = 0.056, *p* = 0.955 for O-S; z = −0.649, *p* = 0.662 for T/O-S; z = 0, *p* = 1.0 for T-O).

**Fig. S3.**
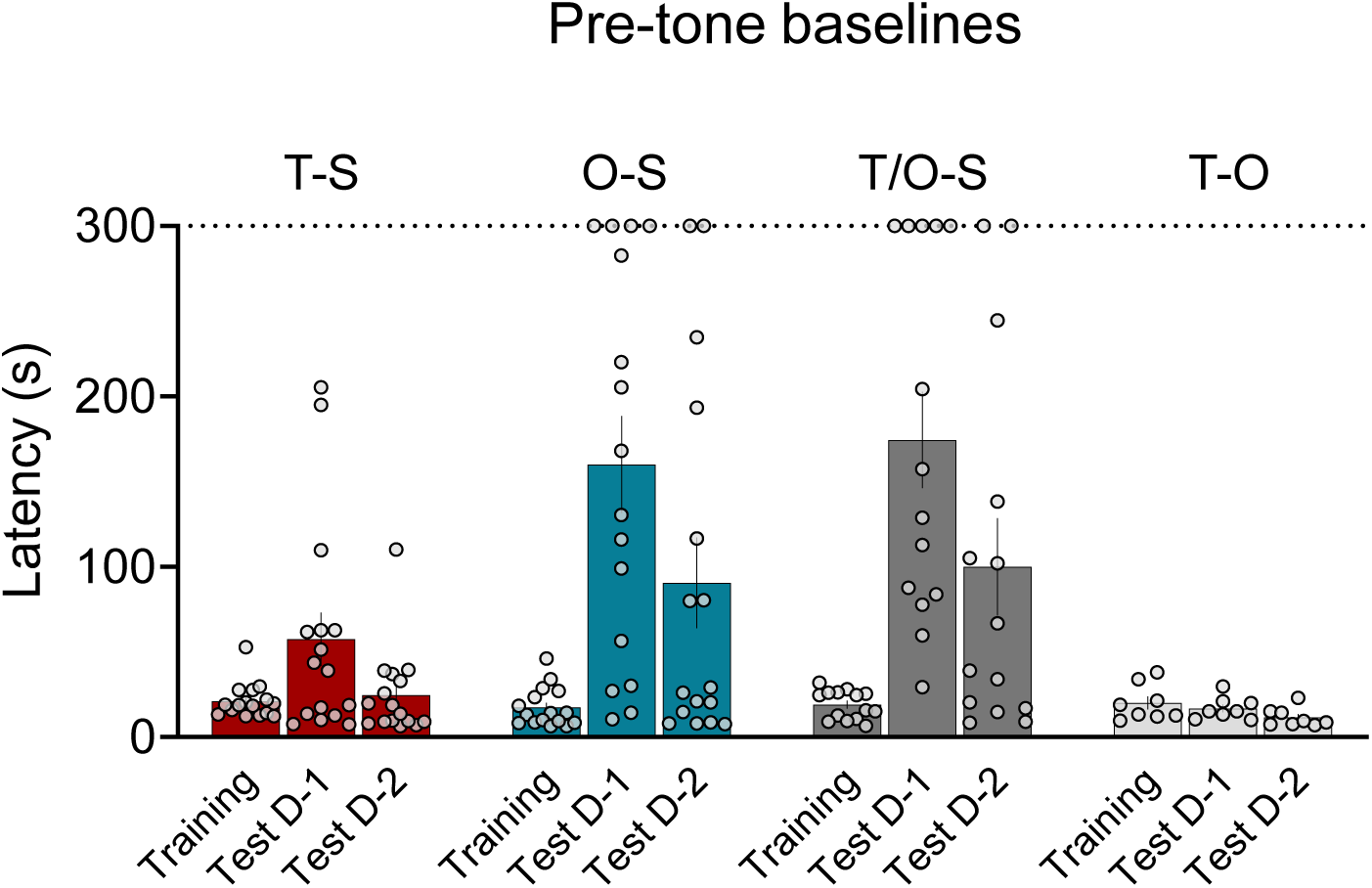
Comparisons of latencies to procure pellets during pre-fear conditioning baseline and pre-tone testing baseline days 1 and 2. The baseline latencies to procure pellets prior to the fear conditioning session (Fig. 2A) were not statistically different from the day 1 (Fig. 3A) and day 2 (Fig. 3C) pre-tone test baseline latencies after the fear conditioning session in both tone-shock (T-S) and tone-owl (T-O) paired animals (Related-samples Wilcoxon signed rank test; Baseline vs. D-1: z = 1.293, *p* = 0.196 for T-S; z = −0.560, *p* = 0.575 for T-O; Baseline vs. D-2: z = −0.155, *p* = 0.877 for T-S; z = −1.82, *p* = 0.069 for T-O). This indicates that neither the tone-shock group nor the tone-owl group showed evidence of contextual fear conditioning.

### Legends for supplementary movies

#### Movie S1

Representative foraging and escape behaviors of a rat presented with an owl-shock pairing. As the animal come near a pellet, it encounters a swooping owl (from behind a black curtain) followed by a dorsal neck/body shock pain. The rat flees to the nest without procuring the pellet.

#### Movie S2

The next day, as the same O-S rat advances towards a pellet, a novel tone is presented for the first time. In response to the tone, the rat promptly flees to the nest without procuring the pellet.

## References

1. Watson JB & Morgan JJB (1917) Emotional reactions and psychological experimentation. Am J Psychol 28(3):163–174.

2. Watson JB & Rayner R (1920) Conditioned emotional reactions. J Exp Psychol 3(3):1–14.

3. LeDoux J (1998) Fear and the brain: where have we been, and where are we going? Biol Psychiatry 44(12):1229–1238.

4. Fendt M & Fanselow MS (1999) The neuroanatomical and neurochemical basis of conditioned fear. Neurosci Biobehav Rev 23(5):743–760.

5. Maren S & Quirk GJ (2004) Neuronal signalling of fear memory. Nat Rev Neurosci 5(11):844–852.

6. Bouton ME, Mineka S, & Barlow DH (2001) A modern learning theory perspective on the etiology of panic disorder. Psychol Rev 108(1):4–32.

7. Kim JJ & Jung MW (2006) Neural circuits and mechanisms involved in Pavlovian fear conditioning: a critical review. Neurosci Biobehav Rev 30(2):188–202.

8. Watson JB (1913) Psychology as the behaviorist views it. Psychological Review 20(2):158–177.

9. Pavlov IP (1927) Conditioned Reflexes: An Investigation of the Physiological Activity of the Cerebral Cortex (Oxford University Press, London) p 142.

10. Guthrie ER (1930) Conditioning as a principle of learning. Psychological Review 37(5):412–428.

11. Kamin LJ (1968) Attention-like processes in classical conditioning in Miami symposium on the prediction of behavior, ed Jones MR (University of Miami Press), pp 9–33.

12. Rescorla RA (1968) Probability of shock in the presence and absence of CS in fear conditioning. J Comp Physiol Psychol 66(1):1–5.

13. Wagner AR, Logan FA, Haberlandt K, & Price T (1968) Stimulus selection in animal discrimination learning. J Exp Psychol 76(2):171–180.

14. rescorla RA & Wagner AR (1972) A theory of Pavlovian conditioning: variations in the effectiveness of reinforcement and nonreinforcement (Appleton-Century-Crofts, New York) p 37.

15. Josselyn SA & Tonegawa S (2020) Memory engrams: Recalling the past and imagining the future. Science 367(6473).

16. Tovote P, Fadok JP, & Luthi A (2015) Neuronal circuits for fear and anxiety. Nat Rev Neurosci 16(6):317–331.

17. Haubensak W, et al. (2010) Genetic dissection of an amygdala microcircuit that gates conditioned fear. Nature 468(7321):270–276.

18. Foa EB & Rothbaum BO (1998) (Guilford Press, New York).

19. Butler AC, Chapman JE, Forman EM, & Beck AT (2006) The empirical status of cognitive-behavioral therapy: a review of meta-analyses. Clin Psychol Rev 26(1):17–31.

20. Delgado MR, Olsson A, & Phelps EA (2006) Extending animal models of fear conditioning to humans. Biol Psychol 73(1):39–48.

21. Mahan AL & Ressler KJ (2012) Fear conditioning, synaptic plasticity and the amygdala: implications for posttraumatic stress disorder. Trends Neurosci 35(1):24–35.

22. Craske MG, et al. (2011) What is an anxiety disorder? Focus 9(3):20.

23. Lima SL & Dill LM (1990) Behavioral Decisions Made under the Risk of Predation - a Review and Prospectus. Canadian Journal of Zoology 68(4):619–640.

24. Bednekoff PA (2007) Foraging in the Face of Danger (University of Chicago Press, Chicago).

25. Stephens DW (2008) Decision ecology: foraging and the ecology of animal decision making. Cogn Affect Behav Neurosci 8(4):475–484.

26. Beckers T, Krypotos AM, Boddez Y, Effting M, & Kindt M (2013) What’s wrong with fear conditioning? Biol Psychol 92(1):90–96.

27. Mobbs D & Kim JJ (2015) Neuroethological studies of fear, anxiety, and risky decision-making in rodents and humans. Curr Opin Behav Sci 5:8–15.

28. Pellman BA & Kim JJ (2016) What Can Ethobehavioral Studies Tell Us about the Brain’s Fear System? Trends Neurosci 39(6):420–431.

29. Bolles RC (1970) Species-Specific Defense Reactions and Avoidance Learning. Psychological Review 77(1):32–48.

30. Choi JS & Kim JJ (2010) Amygdala regulates risk of predation in rats foraging in a dynamic fear environment. Proc Natl Acad Sci U S A 107(50):21773–21777.

31. Zambetti PR, Schuessler BP, & Kim JJ (2019) Sex Differences in Foraging Rats to Naturalistic Aerial Predator Stimuli. iScience 16:442–452.

32. Yilmaz M & Meister M (2013) Rapid innate defensive responses of mice to looming visual stimuli. Curr Biol 23(20):2011–2015.

33. Papes F, Logan DW, & Stowers L (2010) The vomeronasal organ mediates interspecies defensive behaviors through detection of protein pheromone homologs. Cell 141(4):692–703.

34. Tolman EC (1948) Cognitive maps in rats and men. Psychol Rev 55(4):189–208.

35. Wilensky AE, Schafe GE, & LeDoux JE (2000) The amygdala modulates memory consolidation of fear-motivated inhibitory avoidance learning but not classical fear conditioning. J Neurosci 20(18):7059–7066.

36. Lee T & Kim JJ (2004) Differential effects of cerebellar, amygdalar, and hippocampal lesions on classical eyeblink conditioning in rats. J Neurosci 24(13):3242–3250.

37. Stiedl O & Spiess J (1997) Effect of tone-dependent fear conditioning on heart rate and behavior of C57BL/6N mice. Behav Neurosci 111(4):703–711.

38. Guimaraes FS, Hellewell J, Hensman R, Wang M, & Deakin JF (1991) Characterization of a psychophysiological model of classical fear conditioning in healthy volunteers: influence of gender, instruction, personality and placebo. Psychopharmacology (Berl) 104(2):231–236.

39. Rescorla RA (1988) Behavioral studies of Pavlovian conditioning. Annu Rev Neurosci 11:329–352.

40. Thompson RF & Krupa DJ (1994) Organization of memory traces in the mammalian brain. Annu Rev Neurosci 17:519–549.

41. Fanselow MS & Wassum KM (2015) The Origins and Organization of Vertebrate Pavlovian Conditioning. Cold Spring Harb Perspect Biol 8(1):a021717.

42. Lee HJ, Berger SY, Stiedl O, Spiess J, & Kim JJ (2001) Post-training injections of catecholaminergic drugs do not modulate fear conditioning in rats and mice. Neurosci Lett 303(2):123–126.

43. Palgi Y, Gelkopf M, & Berger R (2015) The inoculating role of previous exposure to potentially traumatic life events on coping with prolonged exposure to rocket attacks: A lifespan perspective. Psychiatry Res 227(2-3):296–301.

44. Somer E, et al. (2009) Israeli civilians under heavy bombardment: prediction of the severity of post-traumatic symptoms. Prehosp Disaster Med 24(5):389–394.

45. Alexander BK, Beyerstein BL, Hadaway PF, & Coambs RB (1981) Effect of early and later colony housing on oral ingestion of morphine in rats. Pharmacol Biochem Behav 15(4):571–576.

46. Gage SH & Sumnall HR (2019) Rat Park: How a rat paradise changed the narrative of addiction. Addiction 114(5):917–922.

47. Fanselow MS & Lester LS (1988) A functional behavioristic approach to aversively motivated behavior: Predatory imminence as a determinant of the topography of defensive behavior (Lawrence Erlbaum Associates Inc).

48. Hull CL (1929) A functional interpretation of the conditioned reflex. Psychological Review 36(6):498–511.

49. Lazarus AA (1971) Behavior Therapy and Beyond (McGraw-Hill Companies).

50. Öhman A & Mineka S (2001) Fears, phobias, and preparedness: Toward an evolved module of fear and fear learning. Psychological Review 108(3):483–522.

51. Mathis A, et al. (2018) DeepLabCut: markerless pose estimation of user-defined body parts with deep learning. Nat Neurosci 21(9):1281–1289.

